# Efficient culture of SARS-CoV-2 in human hepatoma cells enhances viability of the virus in human lung cancer cell lines permitting the screening of antiviral compounds

**DOI:** 10.1101/2020.10.04.325316

**Authors:** Santseharay Ramirez, Carlota Fernandez-Antunez, Long V. Pham, Line A. Ryberg, Shan Feng, Martin S. Pedersen, Lotte S. Mikkelsen, Sandrine Belouzard, Jean Dubuisson, Judith M. Gottwein, Ulrik Fahnøe, Jens Bukh

## Abstract

Efforts to mitigate COVID-19 include screening of existing antiviral molecules that could be re-purposed to treat SARS-CoV-2 infections. Although SARS-CoV-2 propagates efficiently in African green monkey kidney (Vero) cells, antivirals such as nucleos(t)ide analogs (nucs) often exhibit decreased activity in these cells due to inefficient metabolization. Limited SARS-CoV-2 replication and propagation occurs in human cells, which are the most relevant testing platforms. By performing serial passages of a SARS-CoV-2 isolate in the human hepatoma cell line clone Huh7.5, we selected viral populations with improved viability in human cells. Culture adaptation led to the emergence of a significant number of high frequency changes (>90% of the viral population) in the region coding for the spike glycoprotein, including a deletion of nine amino acids in the N-terminal domain and 3 amino acid changes (E484D, P812R, and Q954H). We demonstrated that the Huh7.5-adapted virus exhibited a >3-Log_10_ increase in infectivity titers (TCID_50_) in Huh7.5 cells, with titers of ~8 Log_10_TCID_50_/mL, and >2-Log_10_ increase in the human lung cancer cell line Calu-1, with titers of ~6 Log_10_TCID_50_/mL. Culture adaptation in Huh7.5 cells further permitted efficient infection of the otherwise SARS-CoV-2 refractory human lung cancer cell line A549, with titers of ~6 Log_10_TCID_50_/mL. The enhanced ability of the virus to replicate and propagate in human cells permitted screening of a panel of nine nucs, including broad-spectrum compounds. Remdesivir, EIDD-2801 and to a limited extent galidesivir showed antiviral effect across these human cell lines, whereas sofosbuvir, uprifosbuvir, valopicitabine, mericitabine, ribavirin, and favipiravir had no apparent activity.

**Importance:** The cell culture adapted variant of the SARS-CoV-2 virus obtained in the present study, showed significantly enhanced replication and propagation in various human cell lines, including lung derived cells otherwise refractory for infection with the original virus. This SARS-CoV-2 variant will be a valuable tool permitting investigations across human cell types, and studies of identified mutations could contribute to our understanding of viral pathogenesis. In particular, the adapted virus can be a good model for investigations of viral entry and cell tropism for SARS-CoV-2, in which the spike glycoprotein plays a central role. Further, as shown here with the use of remdesivir and EIDD-2801, two nucs with significant inhibitory effect against SARS-CoV-2, large differences in the antiviral activity are observed depending on the cell line. Thus, it is essential to select the most relevant target cells for pre-clinical screenings of antiviral compounds, facilitated by using a virus with broader tropism.

## Introduction

The Severe Acute Respiratory Syndrome Coronavirus 2 (SARS-CoV-2), responsible for COVID-19, was first identified in China in early 2020^1^, but has reached pandemic proportions with over 29 million people infected worldwide by September of 2020^2^. Although a hallmark of COVID-19 is the development of various levels of lung affection from mild upper airway symptoms to life-threatening pneumonia, additional distinctive features of the disease are vascular changes^3^ and numerous extrapulmonary manifestations and systemic complications^4^.

The SARS-CoV-2 virus has been classified within the *Coronaviridae* family, *Betacoronavirus* genus and *Sarbecovirus* subgenus^5^. It is most closely related to SARS-like betacoronavirus of bat origin, but its genomic organization is very similar to the well-characterized SARS-CoV^1^. Its genome consists of a long positive sense, single strand RNA molecule of approximately 30 kb, following the classical genomic organization of viruses belonging to this family, with untranslated regions (UTR) at the 5’ and 3’ ends and numerous open reading frames (ORF) throughout the coding sequences^1^. Separate ORFs encode the structural components, including the spike glycoprotein (S), the envelope (E), the membrane (M), and the nucleocapsid (N), as well as various accessory proteins^1^. The largest ORFs (1a and 1ab) encode the nonstructural proteins of the virus (nsp), responsible for the RNA synthesis machinery^1^. The viral RNA dependent RNA polymerase (nsp12) is an attractive drug target for antiviral therapy. However, identifying nucleos(t)ide analogs (nucs) with anti-coronavirus activity is challenged by the viral nsp14 protein, which has 3’ to 5’ proofreading exoribonuclease capacity, and only a very limited number of molecules have been shown to overcome this activity^6^.

Drug repurposing, compared to de-novo drug discovery, significantly shortens the time and reduces the cost of developing antivirals for emerging pandemic viral diseases. Two of the antiviral molecules that inhibit SARS-CoV-2 and other pandemic coronaviruses are broad-spectrum antivirals remdesivir and β-d-N4-hydroxycytidine (NHC), which were initially part of antiviral discovery programs for the treatment of hepatitis C virus (HCV) infections^7,8^. Remdesivir is now included in standard of care treatment of COVID-19 patients in several countries. It is therefore relevant to continue searching for such compounds, focusing on the molecules that exhibit a good safety profile in humans and developing appropriate models for their pre-clinical screening in cell culture.

SARS-CoV-2 infects the epithelial cells in the upper respiratory tract and in the lungs in humans, however, established cell lines differ in their support for SARS-CoV-2 viral production^9^. Various clones of the African green monkey kidney cell line “Vero” support the highest levels of SARS-CoV-2 replication and propagation, however, not being a human cell line Vero cells show limitations for drug screening, especially for screening of prodrugs that have been designed to be metabolized in human cells^10^. Calu-3, a non-small-cell lung cancer cell line^11^, supports SARS-CoV-2 infection and replication with viral particle production, but at lower levels than those observed in Vero cells^10,12^. Other viruses, such as HCV, also exhibit restricted culture viability, and efficient replication and propagation of HCV is limited to specific clones derived from the human hepatoma cell line (Huh7), including Huh7.5^13,14^.

In this study, we aimed at establishing more efficient and robust platforms for the culture of SARS-CoV-2 in human cells through viral adaptation, permitting basic studies of the virus, including the screening of antiviral compounds, in relevant cell lines.

## Results

### Characterization of a COVID-19 associated SARS-CoV-2 isolate (SARS-CoV-2/human/Denmark/ DK-AHH1/2020) in Vero E6 cells

Vero cells, which permit the isolation of SARS-CoV-2 with high efficiency, were used to obtain the initial virus stock. Vero E6 cells were inoculated with a diluted nasopharyngeal sample from a COVID-19 patient diagnosed with SARS-CoV-2 infection. On day 3 after inoculation, cell cytopathic effect (CPE) was observed, which peaked at day 7. At first, CPE was characterized by the presence of syncytia, followed by the appearance of moderate cell death. Severe CPE with significant cell death was not observed until days 6-7. Culture supernatants were harvested daily during the first 5 days of the experiment and the infectivity titers (expressed as tissue culture infectious dose 50% per milliliter (Log_10_TCID_50_/mL)) were determined using Vero E6 cells and were found to peak at day 4 at 6.4 Log_10_TCID_50_/mL.

The culture supernatant harvested at day 2 post-inoculation was used to perform a first passage infection of naïve Vero E6 cells at a multiplicity of infection (MOI) of 0.02. After passage to naïve cells, CPE was observed at day 2. Two independent second passages were performed using the culture supernatants harvested at days 2 and 3, respectively. CPE was observed promptly after infection, and a large volume of supernatant was harvested at days 2 and 3 from both independent experiments. These supernatants were combined to generate a stock virus (referred to as P2^VeroE6^) that was further characterized and used for the additional experiments described below. The infectivity titer of the P2^VeroE6^ virus stock was 5.5 Log_10_TCID_50_/mL.

We determined the complete SARS-CoV-2 genome sequence (including the 5’ and 3’ UTRs) of the original patient virus and of the P2^VeroE6^ virus as described in Materials and Methods. As shown in Table 1, the isolate SARS-CoV-2/human/Denmark/DK-AHH1/2020, referred to as SARS-CoV-2_DK-AHH1 hereinafter, exhibits only 10 nucleotide differences when compared to the reference sequence of the Wuhan-Hu-1 isolate (genbank NC045512.2). Among these, 6 differences lead to amino acid changes in the nsp2, nsp12, S, ORF3a, and N genomic regions. SARS-CoV-2_DK-AHH1 harbors the high frequency polymorphisms D614G in S, which is now dominant throughout the world and has been linked to increased infectivity in pseudo-typed virus assays in cell culture^15^, and P323L in nsp12, which has not yet been linked to any phenotype. Compared to the original clinical sample, the virus recovered after 2 passages in Vero E6 cells (P2^VeroE6^) was remarkably similar, with only one new change emerging in a small subset of the viral population (C71T in the 5’UTR). Additionally, G22487A (amino acid change E309K in the S-protein), which was the only consensus difference between isolate DK-AHH1 and the reference sequence Wuhan-Hu-1 not fixed (frequency under 99%) in the viral population, further decreased in frequency after passage in Vero E6 cells from 81.8% to 56.9% (Table 1).

**Table 1.** Sequence comparison between the Wuhan-Hu-1 and DK-AHH1 isolates. The near complete SARS-CoV-2 virus genome was sequenced by NGS from 5 overlapping RT-PCR amplicons using a cut-off of 5%, and changes over the cut-off in one sample are shown; to obtain the complete sequence the 5’ and 3’ termini were determined by Sanger sequencing of RT-PCR amplicons. Differences are shown as nucleotide position (number) according to the reference sequence of the Wuhan-Hu-1 isolate (genbank NC045512.2), followed by the specific nucleotide in the reference sequence and in the DK-AHH1 isolate. Original swab refers to the sequence from the patient diagnosed with SARS-CoV-2. The P2^VeroE6^ virus refers to the stock of the second passage in Vero E6 cells. The frequency of the changes in the sequences of the two analyzed viruses (percentage of the reads) is shown in the next two columns. Sequence identity is represented by a dot. Nucleotide sequences were translated *in silico*, and if the nucleotide changes led to a substitution the corresponding amino acid is indicated (original amino acid->new amino acid). Synonymous changes are represented with a dash “-”. The genomic region for each change is also indicated.

| Nucleotide position | Nucleotide in Wuhan-Hu-1 | Nucleotide in DK-AHH1 | Frequency in original swab sample (%) | Frequency in P2<sup>VeroE6</sup> virus (%) | Amino-acid change | Genomic region |
| --- | --- | --- | --- | --- | --- | --- |
| 71 | C | T | . | 6,3 | - | 5'UTR |
| 241 | C | T | 99,7 | 99,6 | - | 5'UTR |
| 1059 | C | T | 99,9 | 99,7 | T->I | nsp2 |
| 3037 | C | T | 99,6 | 99,4 | - | nsp3 |
| 10465 | G | A | 99,8 | 99,7 | - | nsp5 |
| 14408 | C | T | 99,8 | 99,7 | P->L | nsp12 |
| 21646 | C | T | 0,6 | 9,9 | - | S |
| 21742 | C | T | 99,8 | 99,8 | - | S |
| 22487 | G | A | 81,8 | 56,9 | E->K | S |
| 23403 | A | G | 99,9 | 99,7 | D->G | S |
| 25563 | G | T | 99,9 | 99,8 | Q->H | ORF3a |
| 28899 | G | T | 99,9 | 99,5 | R->I | N |

### SARS-CoV-2 can be adapted to efficient growth in human hepatoma cells (cell clone Huh7.5)

We performed inoculations of Huh7.5 cells with the P2^VeroE6^ virus to determine the susceptibility and permissiveness of this cell line. The virus supernatant derived from this first passage was used to inoculate naïve Huh7.5 cells to perform the next passage and successively until passage 6 (Figure 1a). We observed that the emergence of CPE occurred earlier and was more evident as the passage number increased, suggesting viral adaptation. In passage 1 in Huh7.5 cells, CPE was not observed until day 8 post-infection, and the magnitude of the CPE was low at this time, with only minor changes in the morphology of the cells, in contrast to the CPE observed in the passage 6 culture, which displayed massive cell death by day 4 post-infection. The 5^th^ passage in Huh7.5 was performed in big cell flasks, in order to generate a large volume of supernatant that could be used for characterization of the Huh7.5 adapted virus (this virus will be referred to as P5^Huh7.5^ virus).

**Figure 1.**
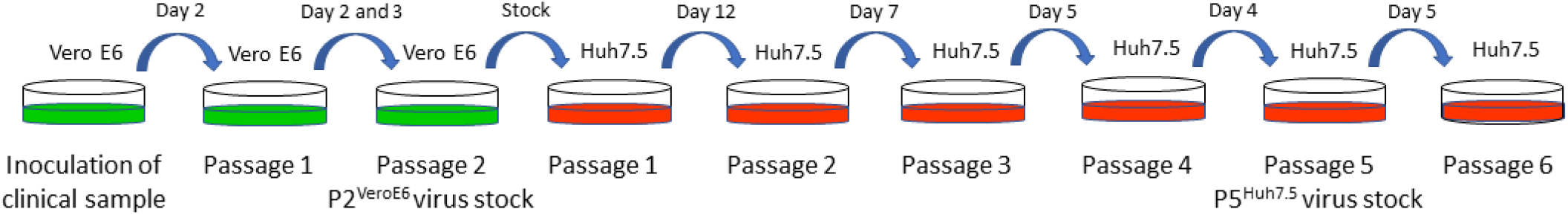
Adaptation of SARS-CoV-2 to efficient growth in Huh7.5 cells. **1a:** schematic overview of the serial passages performed in Vero E6 and Huh7.5 cells. Dishes represent the culture surfaces, green and red color for Vero E6 and Huh7.5 cells, respectively. The harvested day used for passage is indicated above the arrow that symbolizes the transfer of culture supernatant to naïve cells.

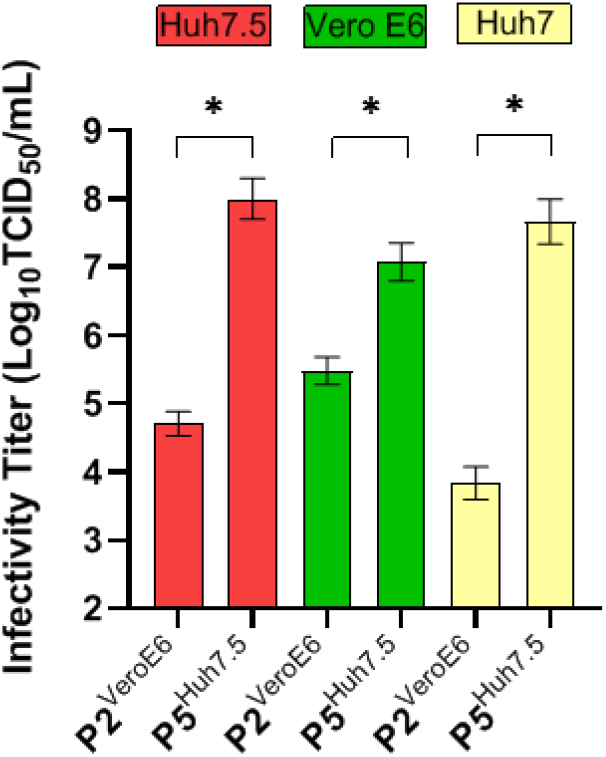
**1b:** comparative infectivity titers of the P2^VeroE6^ and P5^Huh7.5^ viruses in Huh7.5 (red), Vero E6 (green), and Huh7 (yellow) cells. Infectivity titers (Log_10_TCID_50_/mL) are shown on the y-axis. Results are based on several independent experiments: for P2^VeroE6^ and P5^Huh7.5^ in Huh7.5 cells, 6 and 2 independent experiments with 4 replicates each are represented, respectively. In a third independent titration experiment end-point dilution for the P5^Huh7.5^ virus was not achieved (Log_10_TCID_50_/mL> 7) and thus the data was not included in the graph. For the Vero E6 cells, the data presented in the graph corresponds to 5 (P2^VeroE6^) and 4 (P5^Huh7.5^) independent experiments with 4 replicates each. For the parental Huh7 cells, results are based on 3 independent experiments for each virus. Bars represent the mean and standard error of the mean (SEM) of the different independent experiments. Statistical significance (p<0.05, unpaired t-test) is highlighted with an asterisk (*).

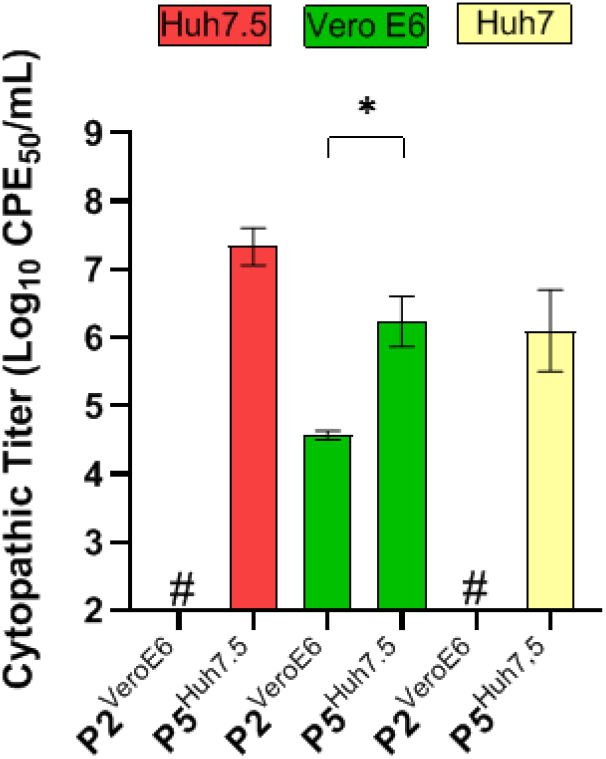
**1c:** comparative cytopathic effect titers of the P2^VeroE6^ and P5^Huh7.5^ viruses in Huh7.5 (red), Vero E6 (green) and Huh7 (yellow) cells. Cytopathic effect titers (Log_10_CPE_50_/mL) are shown on the y-axis. For P2^VeroE6^ and P5^Huh7.5^ in both Huh7.5 and Vero E6 cells, results are based on 3 independent experiments with 4 replicates each. For the P2^VeroE6^ virus in Huh7.5 cells only one experiment yielded a CPE value over the threshold and thus the value was not plotted (depicted with the “#” symbol, threshold calculated as stated in Materials and Methods). In the case of the parental Huh7 cells, results are based on 3 independent experiments as well, and none of the experiments with the P2^VeroE6^ virus yielded values over the assay threshold (#). Bars represent the SEM of the different independent experiments. Statistical significance (p<0.05, unpaired t-test) is highlighted with an asterisk (*).

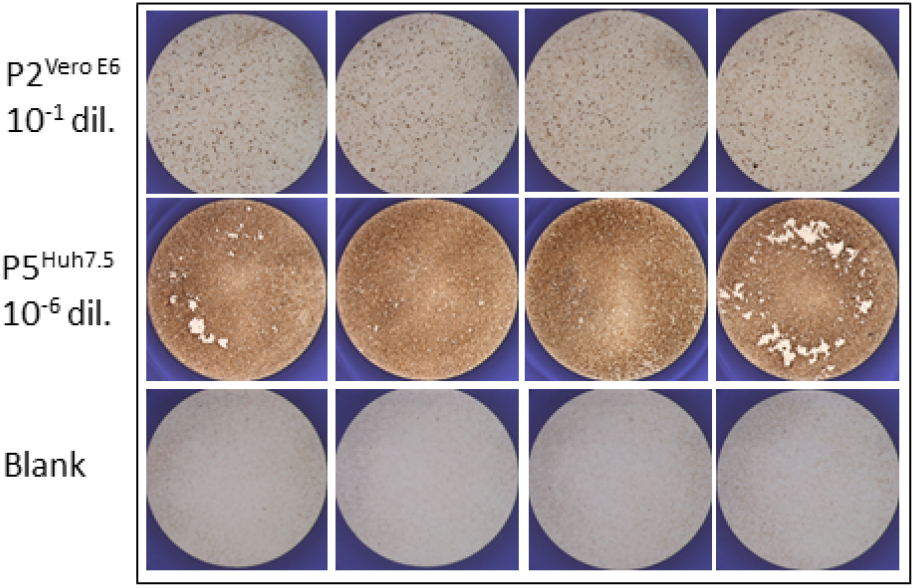
**1d:** visual comparative SARS-CoV-2 antigen staining of both the P2^VeroE6^ and P5^Huh7.5^ viruses infecting Huh7.5 cells and blank (non-infected cells), from a representative TCID_50_ assay. Each picture represents a replicate of infections performed at the indicated dilution (non-cytopathic) for each virus and were obtained after HRP staining with an anti-spike antibody, using the ImmunoSpot series 5 UV analyzer as described in Materials and Methods.

We performed a comparative titration in various cells of the P2^VeroE6^ and the P5^Huh7.5^ viruses (Figure 1b) and found that the infectivity titers in Huh7.5 cells after culture adaptation had increased by more than 3 logs (mean of 4.7 and 8.0 Log_10_TCID_50_/mL, respectively). The Huh7.5 adapted virus also exhibited significantly increased titers in Vero E6 cells (mean of 7.0 Log_10_TCID_50_/mL for the P5^Huh7.5^ virus versus 5.5 Log_10_TCID_50_/mL for the P2^VeroE6^ virus). Interestingly, the original P2^VeroE6^ virus was less viable in the Huh7 parental cell line than in the Huh7.5 clone, however, adaptation to the Huh7.5 clone also led to a significant increase in infectivity titers in Huh7 cells (3.8 and 7.7 Log_10_TCID_50_/mL for the P2^VeroE6^ and P5^Huh7.5^ viruses, respectively).

Visual observations of P5^Huh7.5^ virus infected cultures in the light microscope indicated an increase in CPE. To better quantify this, we performed viral cytopathic effect assays (CPE assays, Figure 1c), in which we detected an evident increase in CPE titers (Log_10_CPE_50_/mL) in all cells. CPE significantly increased from 4.6 to 6.2 Log_10_CPE_50_/mL in Vero E6 cells. In Huh7.5 cells, the P2^VeroE6^ virus was not cytopathic (we obtained a value just above the assay threshold in one of the independent experiments), whereas the P5^Huh7.5^ virus led to high titers of 7.3 Log_10_CPE_50_/mL. For the Huh7 parental cells the P2^VeroE6^ virus was also non-cytopathic, but the adapted P5^Huh7.5^ yielded 6.1 Log_10_CPE_50_/mL.

In addition to the increase in infectivity titers observed after infection with the adapted P5^Huh7.5^ virus in Huh7.5 cells, we also noticed an evident increase in the intensity of the antigen staining (α-spike protein antibody), and in the number of infected cells at non-cytopathic virus dilutions of the P5^Huh7.5^ virus compared to the P2^VeroE6^ virus (Figure 1d). This suggest that the P5^Huh7.5^ virus might both replicate and propagate at higher levels in Huh7.5 cells, as also indicated by the significant increase in CPE titers. In contrast, no such evident differences were observed upon infection of Vero E6 cells with the adapted virus despite the increase in infectivity titers (data not shown).

The near complete genome sequence of the Huh7.5 adapted viruses (P1, P5 and P6) was obtained and analyzed. We found mutations leading to amino acid changes in several nonstructural and structural proteins compared to P2^VeroE6^ (Table 2). The region coding for the spike protein accumulated a significant number of high frequency (>90% of the viral population) changes, including a deletion leading to the removal of nine amino acids in the N-terminal domain (from nt. 21762 to nt. 21788) and 3 non-synonymous mutations: A23014C (E484D aa change according to S-protein specific numbering), C23997G (P812R), and A24424C (Q954H). The deletion and the E484D substitution were already dominant (frequency >90%) after the first passage in Huh7.5 cells (P1^Huh7.5^, Table 2).

**Table 2.** Sequence analysis of the viruses harvested after passage in Huh7.5 cells. The near full-length sequence of the SARS-CoV-2 virus grown in VeroE6 and in Huh7.5 cells (missing the first 30 and the last 65 nucleotides) was obtained by NGS (cut-off 5%). The sequence of the P2^VeroE6^ virus is used as reference and the differences in the Huh7.5 recovered viruses are indicated. “Δ” refers to deletion. The P2^VeroE6^ virus refers to the stock pool of the second passage in Vero E6 cells. For the viruses recovered from Huh7.5 cells, the days used for sequencing were day 12 for P1^Huh7.5^, day 3 (stock) for P5^Huh7.5^ and day 4 for P6^Huh7.5^. For more details refer to Table 1 legend.

| Nucleotide position | P2<sup>VeroE6</sup> | Nucleotide change | P1<sup>Huh7.5</sup> | P5<sup>Huh7.5</sup> | P6<sup>Huh7.5</sup> | Amino-acid change | Genomic region |
| --- | --- | --- | --- | --- | --- | --- | --- |
| 7917 | A | T | 16,7 | 37,5 | 22,2 | E->V | nsp3 |
| 11522 | T | G | 3,1 | 61,9 | 83,7 | F->V | nsp6 |
| 11750 | C | T | 3,9 | 12,3 | 7,0 | L->F | nsp6 |
| 16347 | A | G | . | 12,6 | 19,5 | - | nsp13 |
| 19895 | C | T | . | 30,7 | 33,8 | A->V | nsp15 |
| 20482 | T | C | . | . | 5,0 | S->P | nsp15 |
| 20483 | C | T | . | 0,8 | 5,2 | S->F | nsp15 |
| 21752 | T | A | . | 24,6 | 46,9 | W->R | S |
| 21762...21788 | CTA...GTA | Δ | 100 | 99,9 | 99,9 | Δ9 | S |
| 22110 | A | G | 2,4 | 5,6 | 1,1 | Q->R | S |
| 22264 | C | T | 0,6 | 24,7 | 23,2 | - | S |
| 23014 | A | C | 92,4 | 98,7 | 99,7 | E->D | S |
| 23997 | C | G | . | 92,9 | 99,2 | P->R | S |
| 24424 | A | C | 3,2 | 93,1 | 98,3 | Q->H | S |
| 24983 | T | G | . | 6,7 | 5,9 | L->V | S |
| 26187 | T | C | 82,6 | 85,7 | 81,6 | - | ORF3a |
| 26333 | C | T | 1,0 | 14,8 | 20,9 | T->I | E |
| 28331 | C | T | . | 13,5 | 17,4 | P->S | N |

Outside S, the only other positions in which we found clear indication of evolution at the amino acid level (that is sustained increase in frequency of a residue over the three analyzed Huh7.5 passages) were T11522G (F184V according to nsp6 specific protein number), C19895T (A92V according to nsp15 specific protein number), C26333T (T30I according to E specific protein number), and C28331T (P20S according to N specific protein number) (Table 2).

### Adaptation of SARS-CoV-2 to efficient growth in human hepatoma cells (Huh7.5) leads to robust culture in Calu-1 and A549 lung carcinoma cells

We reasoned that the Huh7.5-adapted viruses might facilitate culture in other human cell lines, including respiratory tract cell lines that would be the most relevant infection model for culture studies. So far, only the Calu-3 cell line has been shown to support, to a limited extent, SARS-CoV-2 productive infections, however, the A549 cell line is refractory to infection^12,16^. To our knowledge, the ability of the Calu-1 cell line to support SARS-CoV-2 replication and propagation has not been previously reported. The Calu-1 and the A549 cell lines are widely available and well-characterized standards among the human lung carcinoma/alveolar cell lines used in cancer research^17,18^. Further, the A549 cell line is also a model for the study of respiratory viruses, such as respiratory syncytial virus and influenza^19,20^.

Compared to the P2^VeroE6^ virus, the P5^Huh7.5^ virus exhibited a significant increase in the ability to infect Calu-1 cells with >2-log increase in infectivity titers. For the original P2^VeroE6^ virus, observed titers in Calu-1 were 3.5 Log_10_TCID_50_/mL, which increased to 6.0 Log_10_TCID_50_/mL for the P5^Huh7.5^ virus (Figure 2a). Surprisingly, the P5^Huh7.5^ virus was able to efficiently infect A549 cells, with titers of 6.0 Log_10_TCID_50_/mL; the A549 cells could not be infected with the P2^VeroE6^ virus (Figure 2a). Albeit at lower levels than in Huh7.5, we could also detect CPE in Calu-1 and A549 cells infected with the P5^Huh7.5^ virus, for which no CPE was observed with the original virus (Figure 2b), indicating enhanced replication.

**Figure 2.**
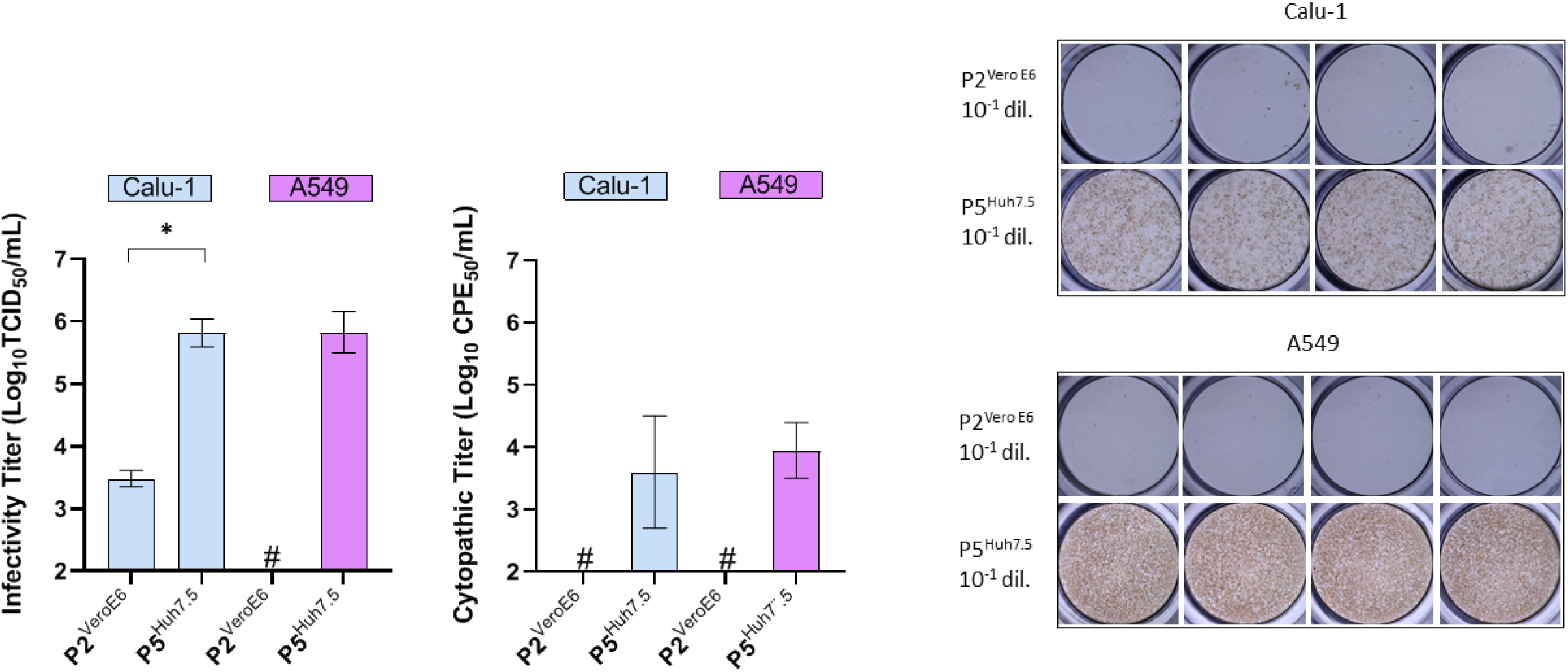
Adaptation of SARS-CoV-2 to Huh7.5 cells permits culture of human lung carcinoma cells Calu-1 and A549. **2a:** comparative infectivity titers of the P2^VeroE6^ and the P5^Huh7.5^ viruses in Calu-1 (blue) and A549 (magenta) cells. Infectivity titers (Log_10_TCID_50_/mL) are shown on the y-axis. For each cell line, results are based on 3 independent experiments with 4 replicates each. For the P2^VeroE6^ virus in A549 cells none of the experiments yielded detectable titers (#). Bars represent the mean and SEM of the different independent experiments. Statistical significance (p<0.05, unpaired t-test) is highlighted with an asterisk (*). **2b:** comparative cytopathic effect of the P2^VeroE6^ and the P5^Huh7.5^ viruses in Calu-1 (blue) and A549 (magenta) cells. Cytopathic effect titers (Log_10_CPE_50_/mL) are shown on the y-axis. Results are based on two independent experiments with 4 replicates each. CPE was under the detection limit in the two experiments with the P2^VeroE6^ virus in both cell lines (#). **2c:** visual comparative SARS-CoV-2 antigen staining of P2^VeroE6^ and P5^Huh7.5^ in Calu-1 (upper panel) and A549 (lower panel), from a representative TCID_50_ assay. Pictures represent 4 replicates of infections performed with the indicated virus dilution (from a representative TCID_50_ assay) and were obtained after HRP staining with an anti-spike antibody, using the ImmunoSpot series 5 UV analyzer as described in Materials and Methods.

In addition to the increase in infectivity and CPE titers observed after infection of the Calu-1 and A549 cells with the P5^Huh7.5^ compared to the P2^VeroE6^ virus, we also detected an increment in the intensity of the antigen staining (α-spike protein antibody) and in the number of infected cells upon inoculation with the same dilution of these viruses (Figure 2c). Thus, and in accordance with our observations in Huh7.5 cells, both better replication and propagation was achieved with the P5^Huh7.5^ virus.

### Improved activity of remdesivir against SARS-CoV-2 in human Huh7.5 cells

Remdesivir was previously reported to inhibit SARS-CoV-2 in Vero^21–23^ and human lung cells (Calu-3)^10^. Here, we found EC_50_ values of 1.5 μM in the concentration-response assays with the P2^VeroE6^ virus in Vero E6 cells (Figure 3a). Further we demonstrated that this drug displayed ~50-fold higher activity against the same virus (P2^VeroE6^) in Huh7.5 cells (EC_50_ of 0.03 μM, Figure 3b). For comparison, we found that remdesivir is a very potent inhibitor of different genotypes of HCV in Huh7.5 cells, with EC_50_ values between 0.08-0.19 μM (Figure 3c). Lastly, we observed that the Huh7.5 adapted virus (P5^Huh7.5^) yielded similar EC_50_ values for remdesivir as the P2^VeroE6^ virus when tested in Huh7.5 cells (EC_50_ values of 0.05 μM and 0.03 μM, respectively) (Table 3 and Figure 3b). Thus, the adapted P5^Huh7.5^ virus represents a useful virus for the screening of SARS-CoV-2 polymerase inhibitors in human hepatoma cells.

**Table 3.** Anti-SARS-CoV-2 activity of a panel of nucleos(t)ide analogs in different human cells. For each compound, the antiviral activity in Huh7.5, Calu-1 or A549 cells is indicated by EC_50_ values (μM). These values were inferred from concentration-response curves as shown in Figure 4. All compounds were tested at non-cytotoxic concentrations as described in Materials and Methods. For remdesivir the maximum concentration tested was 10 μM. For EID-2801 the maximum concentration tested was 50 μM. “>50” indicates that the maximum concentration tested was 50 μM and that no viral inhibition tending towards or reaching 50% was observed at this concentration. The maximum concentration tested for galidesivir was 100 μM (since clear inhibitory effects were observed at 50 μM and 100 μM was non-cytotoxic).

|  | EC<sub>50</sub> (μM) |  |  |
| --- | --- | --- | --- |
|  | Huh7.5 | Calu-1 | A549 |
| Remdesivir | 0.05 | 0.29 | 0.31 |
| EIDD-2801 | 8.5 | 2.7 | 1.3 |
| Galidesivir | 48 | 58 | 24 |
| Sofosbuvir | >50 | >50 | >50 |
| Uprifosbuvir | >50 | >50 | >50 |
| Valopicitabine | >50 | >50 | >50 |
| Mericitabine | >50 | >50 | >50 |
| Ribavirin | >50 | >50 | >50 |
| Favipiravir | >50 | >50 | >50 |

**Figure 3.**
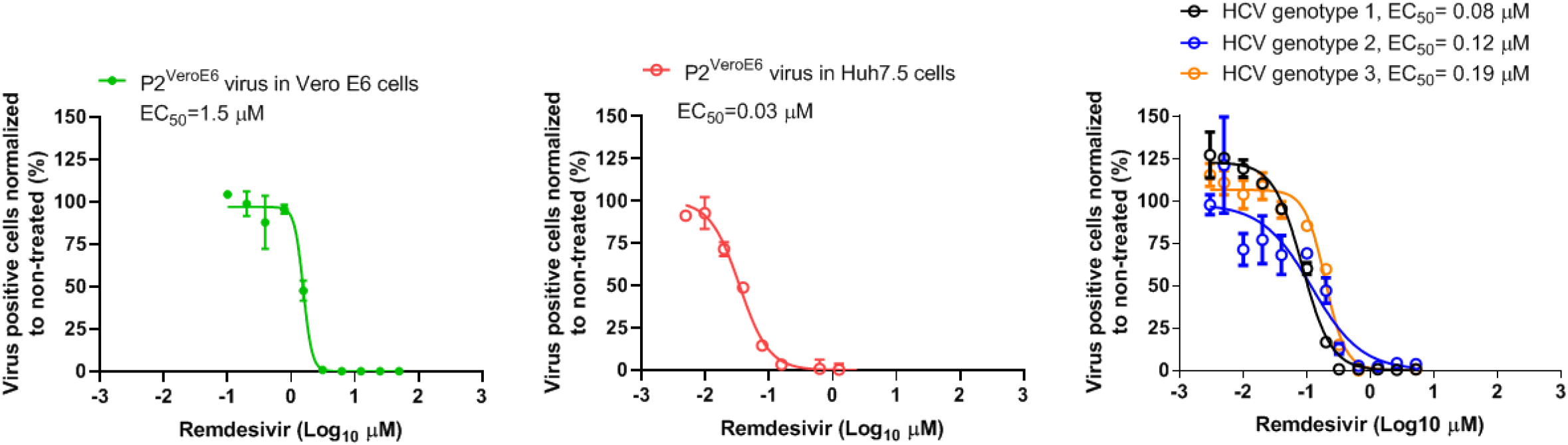
Antiviral activity of remdesivir against SARS-CoV-2 in Vero E6 and Huh7.5 cells and comparison with its activity against HCV genotypes 1-3 in Huh7.5 cells. **3a and b:** antiviral activity of remdesivir against the P2^VeroE6^ virus in Vero E6 cells (3a) and in Huh7.5 cells (3b). The graph shows the non-linear regression curve of the number of SARS-CoV-2 infected cells normalized to non-treated controls (y-axis) of triplicate wells of infected cells treated with different remdesivir concentrations (Log_10_, x-axis). The dots represent the mean, and SEM are shown as lines. EC_50_ values (drug concentration unit is μM) inferred from the regression are shown as well. **3c:** antiviral activity of remdesivir against different genotypes of HCV (genotype 1, strain TNcc in black, genotype 2, strain J6/JFH1 in blue and genotype 3, strain DBN3acc in orange) in Huh7.5 cells performed as previously described for sofosbuvir studies^45–48^.

### Screening of the antiviral activity of a panel of nucs in human cells, using adapted SARS-CoV-2

The adapted P5^Huh7.5^ virus permitted drug testing in Huh7.5, Calu-1 and A549 cells. We focused our screen on nucs previously shown to have antiviral effect against HCV, including sofosbuvir, a pangenotypic HCV drug used in the clinic^24^, but we also included broader-spectrum molecules such as galidesivir, favipiravir and ribavirin.

Among the nucs tested only remdesivir and EIDD-2801 displayed significant effect across the human cells (Table 3), as previously described^25^. Remdesivir was most active in Huh7.5 cells, with ~6-fold lower EC_50_ values when compared to Calu-1 and A549 cells (Figure 4a). Despite being less active in lung carcinoma than in hepatoma cells, remdesivir was still more active in lung carcinoma Calu-1 and A549 than in Vero E6 cells (about 5-fold lower EC_50_ compared to the P2^VeroE6^ virus). The opposite was observed for EIDD-2801 (Figure 4b), which was more active in A549 and Calu-1 cells (6- and 3-fold more active with EC_50_ of 1.3 μM and 2.7 μM, respectively) than in Huh7.5 cells (8.5 μM). Finally, galidesivir exhibited limited activity (Figure 4c) with relatively high EC_50_ (>20 μM); the best inhibitory effect was observed in A549 cells (EC_50_ of 24 μM). Other nucs, including sofosbuvir, had no apparent activity (EC_50_ >50 μM) in these human cell lines with our experimental conditions (Table 3).

**Figure 4.**
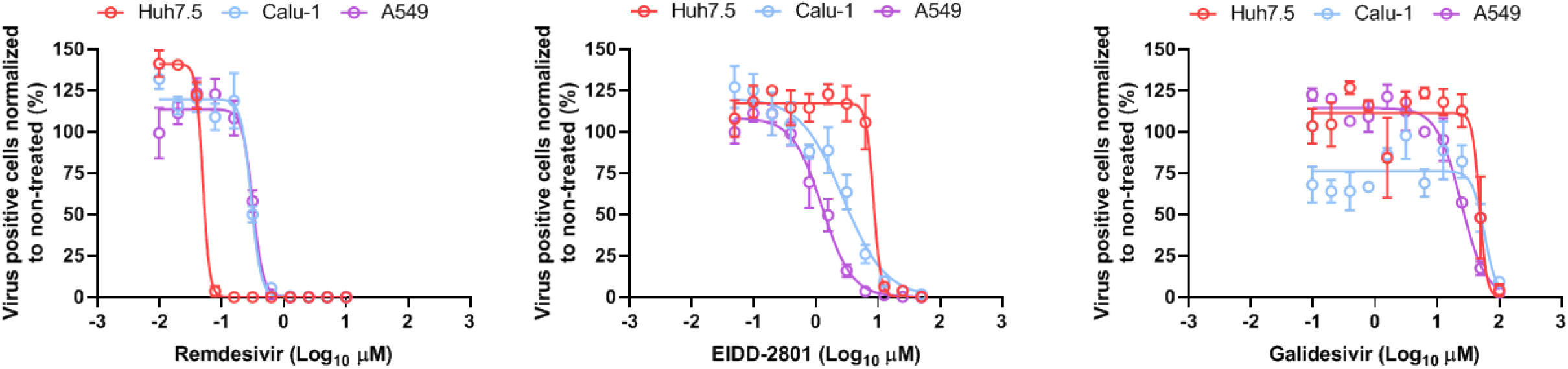
Comparative antiviral activity of remdesivir, EIDD-2801 and galidesivir against SARS-CoV-2_DK-AAH1 in different human cell lines. **4a, b and c:** comparative antiviral activity of remdesivir (4a), EIDD-2801(4b) and galidesivir (4c) against the P5^Huh7.5^ virus in 3 different human cell lines (Huh7.5, red; Calu-1, blue; and A549, magenta). The graph shows the non-linear regression curve of the number of SARS-CoV-2 infected cells normalized to non-treated controls (y-axis) of triplicate wells of infected cells treated with different drug concentrations (Log_10_, x-axis). The dots represent the mean, and SEM are shown as lines. EC_50_ values can be found in Table 3.

## Discussion

In this study, we performed isolation of SARS-CoV-2 (isolate DK-AHH1) in Vero E6 cells and subsequent serial passage of the virus in human hepatoma cells (Huh7.5 clone) that led to adaptation of the virus with increased infectivity and cytopathic effect titers in both Vero E6 and Huh7.5 cells. Most importantly, human hepatoma cell culture adaptation significantly increased the viral viability in lung carcinoma cells Calu-1 and A549, as well as in the parental Huh7 cell line, which do not efficiently support replication and propagation of the original SARS-CoV-2 virus. Thus, the adapted virus might permit culture across additional human cell lines, a proposition that should be the topic of future investigations. We demonstrated that the adapted virus is an efficient tool for the screening of putative SARS-CoV-2 antiviral compounds in human cells. Active molecules, such as remdesivir and EIDD-2801, exhibited increased activity in human cells when compared to Vero E6, highlighting the importance of testing nucs in the appropriate target cells.

The Huh7.5 cell line was originally selected to permit higher replication levels of HCV subgenomic replicons^13^ and it has been of fundamental importance for the development of efficient full-length cell culture systems for HCV^14^. Compared to the parental Huh7, the Huh7.5 clone exhibits a mutation that inactivates the retinoic acid-inducible gene I (RIG-I), an interferon-inducible cellular helicase involved in the type-1 interferon response, one of the features that has been correlated to increased permissiveness to HCV replication^26^. Interestingly, Huh7 cells are highly susceptible and permissive to SARS-CoV, which induces lytic and productive infections^27^. A recent study demonstrated that SARS-CoV-2 is more sensitive to type-1 interferons than SARS-CoV^28^. Therefore decreased type-1 interferon responses might be a key feature for enhancing the cell culture viability of SARS-CoV-2, correlating with the high permissiveness of the Vero cell lines, which lack genes encoding type-1 interferons^29^.

Adaptation of SARS-CoV-2 to Huh7.5 cells caused an accumulation of substitutions in the S genomic region, which might be the most important contributors to the significant increase in the viability of the virus in lung carcinoma cells. The first complete change (present at frequencies of >99% after the first passage) was the deletion of a loop (IHVSGTNGT) in the N-terminal domain of S. This deletion leads to the removal of amino acid N74, which is N-glycosylated^30^; this loop is not present in SARS-CoV, but it is found in MERS-CoV (Figure 5a,b). However, it is unclear how this change could contribute to enhanced viability in culture. Another change that emerged early in most viruses was E484D, which is located in the receptor binding motif of the receptor binding domain. Noticeably, an *in silico* study of E484D predicted higher ACE2 binding affinity that could render a more infectious SARS-CoV-2 virus^31^.

**Figure 5.**
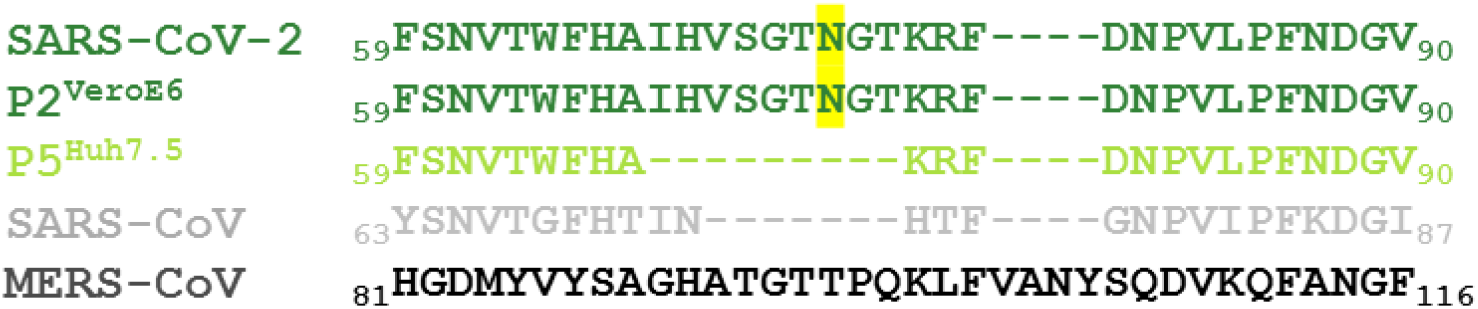
Structural overview of changes found in the S-protein of SARS-CoV-2 after culture adaptation in Huh7.5 cells. Multiple alignment of the partial S-protein sequences of SARS-CoV-2 (GenBank accession number MN908947), P2^VeroE6^ virus, P5^Huh7.5^ virus, SARS-CoV (GenBank accession number AY278741) and MERS-CoV (GenBank accession number JX869059). The alignment was carried out using the MUSCLE software^63^. **5a:** Alignment of the area containing the N-terminal 9 amino acid deletion in Huh7.5 adapted viruses, which corresponds to the protein specific positions indicated with numbers. The SARS-CoV-2 sequences are shown in green with the N74 glycosylation site highlighted in yellow, the SARS-CoV sequence in grey and the MERS-CoV sequence in black.

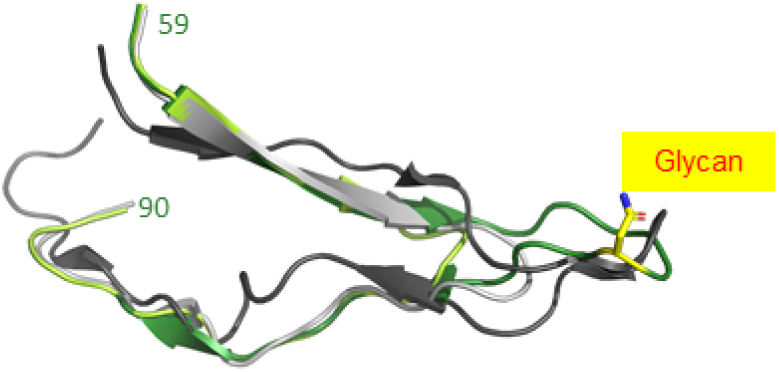
**5b:** Structural alignments of SARS-CoV and MERS-CoV spike proteins (PDB entries 5X58^64^ and 6Q04^65^, respectively) to the SARS-CoV-2 spike protein (PDB entry 7JJI^66^) using PyMOL^67^ with the same parts of the sequences shown in *5a*. The same color coding as in *5a* is used. N74 is shown as sticks in SARS-CoV-2 and an attached glycan is illustrated schematically. The P5^Huh7.5^ structure was generated from the SARS-CoV-2 structure by introducing the deletion and modelling the loop closure (“HAKRFD”) with ModLoop^68^.

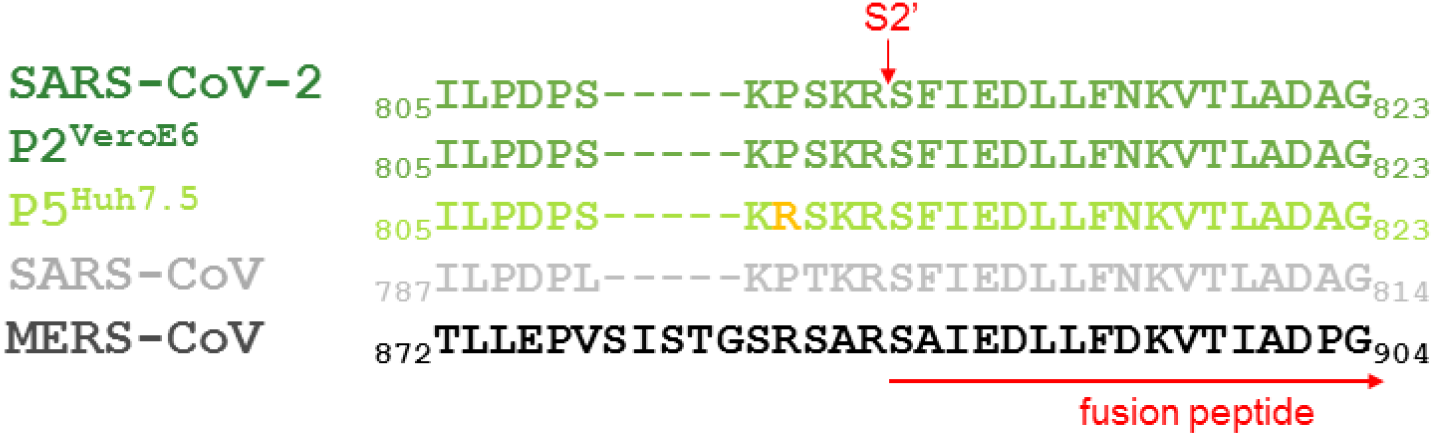
**5c:** Alignment of the area around the S2’ cleavage site (indicated with a red arrow). The observed P812R mutation is highlighted in orange.

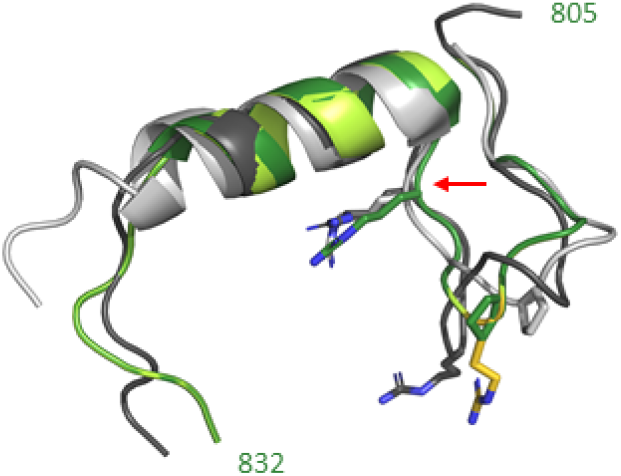
**5d:** Structural alignments with the same parts of the sequences as shown in *5c*. Structures were generated as explained in *5b* using the same PDB entries except for the SARS-CoV spike protein (PDB entry 5XLR^69^). Residues that align with P812 and R815 in SARS-CoV-2 are represented as sticks. A part of the MERS-CoV sequence (”SISTGSRS”) was modelled using ModLoop^68^, as the residues were missing in the experimental structure. The S2’ cleavage site is indicated with a red arrow.

P812R emerged only in the latest passages, when the virus exhibited maximum infectivity. P812 is positioned near the S2’ cleavage site. The proline is also present in SARS-CoV, however, at that position, R is present in MERS-CoV (Figure 5c,d). P812R changes the sequence from “PSKR” to “RSKR”, which corresponds to the furin consensus cleavage motif (“RX[K/R]R”)^32,33^, thus a putative second furin cleavage site could have emerged at the S2’site. Whether the emergence of a new furin cleavage motif nearby the S2’ plays a role in culture adaptation by increasing membrane fusion and therefore infectivity as seen in SARS-CoV, warrants further investigation^34,35^. Similarly, it was found that cell culture adaptation (Vero cells) of the coronavirus infectious bronchitis virus (Beaudette strain) led to the acquisition of a mutation in the S-protein creating a novel furin site downstream of the S1/S2 site that was implicated in the entry and syncytium formation in Vero cells^36^. On the other hand, deletions in the S1-S2 furin cleavage site have been found during culture adaptation in Vero E6 cells^23,37^. Finally, Q954H was present at low frequency in P1 but increased significantly in the last passages, consistent with the maximum increase in viral infectivity. Residue Q954 locates within the S2 subunit, which undergoes conformational rearrangements from prefusion to post fusion states. Specifically, this position is located within the heptad repeat 1 (HR1), which is part of the fusion-active core structure^38^.

As described here by us and elsewhere by others, the A549 cell line is refractory to infection with SARS-CoV-2^16^. For SARS-CoV, it was initially shown that low susceptibility of A549 correlated to a lack of ACE2 expression^39^ and that exogeneous expression supported enhanced SARS-CoV replication. However, natural ACE2 expression in A549 cells was reported by other groups^40,41^.

Another essential host entry factor for SARS-CoV-2 is the transmembrane protease, serine 2 (TMPRSS2)^32^. Vero E6 expressing TMPRSS2 permit enhanced isolation of SARS-CoV-2^12^. TMPRSS2 is also essential for entry of SARS-CoV, specifically the isoform 1 of the protein, directly linked to the activation of the spike protein^42^. This isoform was found to be expressed in virus target cells (such as Calu-3), but was lacking in the A549 cells^42^. Interestingly, TMPRRS2 was not detected in Huh7 cells^43^ either, and thus this could be one of the contributors to the poor virus viability in A549 and in Huh7 cells. It is tempting to speculate that a possible explanation for the significant enhanced viability of the Huh7.5 adapted virus in A549 and Huh7 cells, could relate to the acquisition of the furin cleavage site by substitution P812R, which could compensate for a putative low activation of the spike due to inadequate TMPRRS2 expression. This finding merit further investigations.

The efficient culture of SARS-CoV-2 permitted the screening of a class of polymerase inhibitors (nucs) in relevant cell culture models. As previously described by others, we found that nucs remdesivir and EIDD-2801 exhibited increased activity in human cells compared to Vero cells^10,25^. For remdesivir, maximum activity was observed when using human hepatoma cells, as expected since this compound originated from the HCV antiviral program, and like sofosbuvir, is masked by a McGuigan prodrug moiety, leading to a significant accumulation of the active compound in the liver^44^. In the present study, we also show proof-of-concept of the potent inhibitory effect of remdesivir against recombinants of HCV genotypes 1, 2 and 3 that recapitulate the entire viral life cycle, with EC_50_ values lower than those routinely obtained with sofosbuvir^45–48^. For EIDD-2801, maximal activity was observed in lung cells, in agreement with previous studies showing high activity of the drug in human airway epithelia infected with influenza virus^49^. The EC_50_ values of remdesivir in Vero E6 and Calu-1 cells obtained in this study were very similar to those reported before for Vero and Calu-3 cells, respectively^10^. However, for EIDD-2801 (oral bioavailable prodrug of NHC) the EC_50_ values in Calu-1 and A549 cells were higher than previously reported NHC values in Calu-3^25^, which could be explained by the use of the prodrug in our study.

Our investigated panel of nucs included molecules that had been previously shown to inhibit the virus (remdesivir and EIDD-2801), but also compounds that had been proposed as drug candidates. Strikingly no other analogs than remdesivir and EIDD-2801 showed a significant inhibitory effect in our experimental conditions, including sofosbuvir, which had been the subject of extensive *in silico* investigations^50^ and was found to cause chain termination of the SARS-CoV-2 polymerase *in vitro*^51^. It is important to acknowledge that studies only assessing incorporation and chain termination of nucleotide analogs as antiviral strategies against coronaviruses lack a complete overview of the structural requirements for viral replication, including the role of the nsp14 exonuclease in excising these molecules, which remdesivir overcomes^52^. Favipiravir, a broad-spectrum antiviral with similar mechanism of action to EIDD-2801, exhibited no antiviral effect in our experimental conditions, but contrarily showed anecdotal activity in patients^53^. However proper randomized trials are needed to confirm these findings.

The broad-spectrum nuc galidesivir^54^ exhibited a detectable inhibitory effect against SARS-CoV-2, but significantly lower activity when compared to remdesivir and EIDD-2801. In light of the detectable inhibitory effect, it could be relevant to test the efficacy of this drug in primary airway cultures and in longer-term treatment assays where multiple doses of the compound are added into the culture to further elucidate the potential antiviral use of this compound.

We demonstrated that the activity of remdesivir in Huh7.5 cells was very similar between the P2^VeroE6^ and the adapted P5^Huh7.5^ virus, despite the later exhibiting multiple changes in the genome. Since these changes concentrated in the S-protein and not in the nsp12 protein, which is the main target of nucs, this virus represents an excellent tool to study this drug class in human cells. However, the mutations present in the adapted virus could potentially interfere with entry processes and therefore this experimental system might not be an optimal tool for the screening of entry/fusion inhibitors.

It is important to continue evaluating antiviral strategies against SARS-CoV-2, that could also be applied to other pandemic coronaviruses. Our cell culture adapted virus with increased viability in human cells could contribute to a more efficient and relevant screening of candidate compounds inhibiting viral replication. The cell-culture adaptation strategy shown in this study can represent a venue for the efficient culture of other coronaviruses with restricted viability in human cell lines.

## Materials and Methods

### Cell culture experiments

All experiments in this study were performed with a novel isolate of the SARS-CoV-2 virus (SARS-CoV-2/human/Denmark/DK-AHH1/2020). Initial culture of SARS-CoV-2_DK-AHH1 was performed in African green monkey kidney cells (Vero E6). Residual nasopharyngeal sample material was obtained after clinical diagnosis with COVID-19 (SARS-CoV-2 Ct value of 14 in RT-PCR assay^55^) and irreversible anonymization. For inoculation, 30 μL of the diluted sample were added into 100,000 Vero E6 cells, seeded in 12-well plates, 24 hour prior, at a final volume of 1 mL. For the initial inoculation experiment, Vero E6 cells were cultured in DMEM (high glucose, GlutaMAX and pyruvate, Invitrogen) supplemented with 10% fetal bovine serum (Sigma) and Antibiotic-Antimycotic (10,000 units/mL of penicillin, 10,000 μg/mL of streptomycin, and 25 μg/mL of Amphotericin B, Gibco) and kept at 37°C in a humidified incubator with 5% CO_2_. Afterwards, all experiments in Vero E6 were performed with media supplemented only with FBS 10%, 10,000 units penicillin and 10 mg streptomycin/mL (Sigma). The culture was visually inspected under an inverted light microscope and supernatant was harvested daily and stored at −80°C. A first viral passage from the original inoculated culture was performed with 0.5 mL of harvested supernatant from day 2 post-inoculation in naïve Vero E6 cells in a T-25 flask with 10^6^ cells, with a final volume of 4 mL (approximate MOI of 0.02). Two independent second passages were performed with supernatants harvested at passage 1 day 2 (approximate MOI of 0.9) and day 3 (MOI not determined), respectively. For that purpose, 1.5 mL of first passage supernatant was used to infect 5×10^6^ cells (final volume of 30 mL). A large viral stock was prepared by pooling 30 mL of filtered supernatant (0.45 μM filter, Sartorius) from the samples harvested at days 2 and 3 from both second passages (total of 120 mL), aliquoted, and stored at −80°C.

Human hepatoma cells (Huh7 and Huh7.5 cells) and Calu-1 were cultured in DMEM (high glucose, GlutaMAX and pyruvate, Invitrogen ThermoFisher) supplemented with 10% fetal bovine serum (Sigma), 10,000 units penicillin and 10 mg streptomycin/mL (Sigma), and kept at 37°C in a humidified incubator with 5% CO_2_. A549 cells were cultured in the same conditions but with Ham’s F-12K (Kaighn’s) Medium (Gibco, ThermoFisher) supplemented with the same 10% fetal bovine serum and antibiotics. Culture of HCV was performed as previously described^14,45,46^.

### Serial passage of the SARS-CoV-2 virus in Huh7.5 cells

A total of six passages were conducted in Huh7.5 cells. Passages were performed by inoculating 10^6^ naïve cells (seeded in a T-25 flask) with a 1/10 dilution of the P2^VeroE6^ virus (P1) or with 0,5 mL of supernatant for subsequent passages. The fifth passage was carried out in T-175 flasks (approximately 5 million cells seeded), for production of a large volume to generate a viral stock, which was used for the subsequent assays.

### Determination of viral titers

Infectivity titers were expressed as 50% tissue culture infectious dose per mL (TCID_50_/mL). TCID_50_ assays were performed in 96-well plates, by infecting naïve cells with 100 μL of 10-fold serially diluted virus containing supernatants, in quadruplicates, followed by immunostaining of the SARS-CoV-2 spike protein (as described below) at 72-hr post infection. Presence or absence of infected cells in each replicate was scored and used to determine TCID_50_/mL values based on calculations obtained by the Reed and Muench method (as described in Fields Virology, 5^th^ edition^56^).

A CPE assay to determine the cytopathic effect 50% titer (CPE_50_/mL) was developed with similar conditions as those for the infectivity assay with infections of naïve cells with 100 μL of 10-fold dilutions of virus containing supernatants, in quadruplicates. CPE was assessed 72-hr post infection with the Viral ToxGlo Assay (Promega) following the manufacture’s instructions. Relative light units (RLU) obtained for each infected well was normalized to the RLU of non-infected controls (100% cell normality or cell viability), values under 90% were considered positive for CPE, and the Reed and Muench method was used to calculate a Log_10_ CPE_50_/mL titer. The assay was validated by correlating titer values obtained with the Viral ToxGlo Assay with visual inspections of the cells under a light microscope in multiple assays.

All graphical representations and statistical analysis were performed using GraphPad Prism 8.

### Sequencing of SARS-CoV-2 viruses

The viral sequencing of SARS-CoV-2 recovered from the original clinical specimen and from all cell-culture derived supernatants was performed in an identical manner with next-generation sequencing of near-full-length genomes from 5 overlapping-amplicons in each pool. The methodologies for RNA extraction, generation of cDNA by reverse-transcription (RT) and amplification of these overlapping amplicons were adapted from existing protocols for the amplification of the complete open-reading frame of HCV^57^. Specific SARS-CoV-2 primers can be found in Table 4. Determination of the 5’UTR of the original clinical specimen and of the cell culture recovered P2^VeroE6^ viruses was determined using a template-switching RT method and reagents from New England Biolabs and following guidelines. Gene-specific RT primers can be found in Table 4. The 3’UTR was determined by RT-PCR with an oligo-dT anchor primer (Sigma) as described^58^ with primers found in Table 4. NGS analysis was performed as described previously^59^ with minor modifications. In short, reads were trimmed from PCR primer sequences at the 5’ end by Cutadapt^60^ to remove the bias of the 5 overlapping amplicons. Next, reads were mapped to the Wuhan-Hu-1 reference sequence. Subsequently, consensus and low frequency SNPs and indel calling was performed.

**Table 4.**
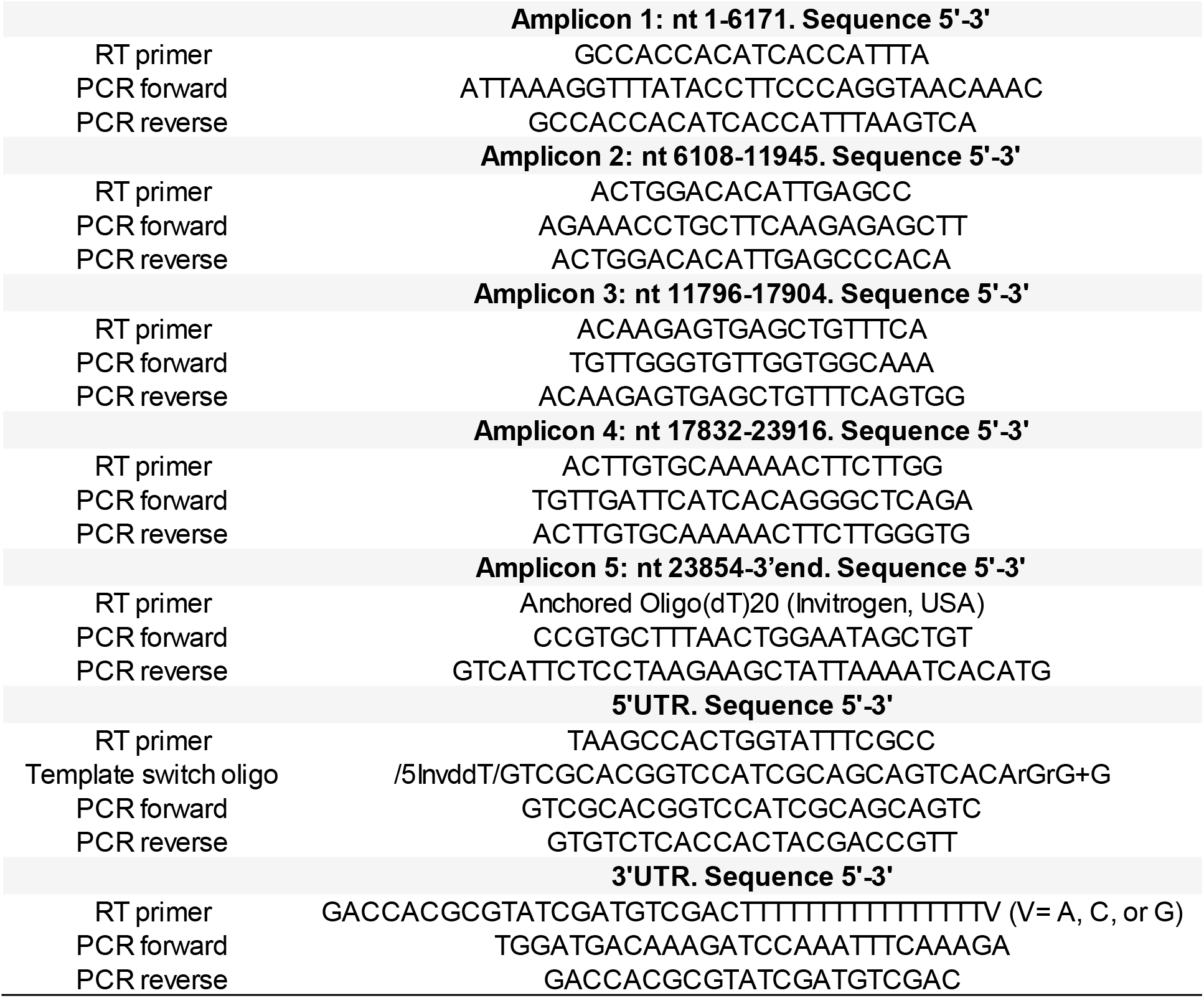
Primers used for SARS-CoV-2 genome amplification procedures. RT (reverse-transcription) and PCR primers (forward and reverse) for each amplicon covering the near full-length PCR strategy (a single RT reaction was performed by pooling all the 5 RT primers, which served as template for each of the independent PCR reactions generating the 5 overlapping amplicons) and the ends of the untranslated regions (UTR) are shown in the table. Nucleotide numbering according to the Wuhan-Hu-1 isolate (genbank NC_045512.2). RT and PCR primers for the determination of the 3’UTR were obtained from Sigma. The remaining primers were purchased from TAG Copenhagen.

### Immunostaining of SARS-CoV-2 infected cell cultures

Following different assays, 96-well plates with confluent cell layers were fixed and virus was inactivated by immersion in 100% methanol (MERCK) for 20 min. Cells were washed twice with PBST (PBS containing 0.1% Tween-20), followed by incubation with 3% H_2_O_2_ for 10 min to block endogenous peroxidase activity and washed twice with PBST. Cells were then incubated with primary antibody SARS-CoV-2 spike chimeric monoclonal antibody (Sino Biological #40150-D004) diluted 1:5000 in PBS with 1% bovine serum albumin and 0.2% skim milk powder (PBSK) overnight at 4 ℃. Afterwards, cells were washed twice with PBST followed by 1-hour incubation with secondary antibody F(ab’)2-Goat anti-Human IgG Fc Cross-Adsorbed Secondary Antibody, HRP (Invitrogen #A24476) diluted 1:2000 in PBSK. Cells were washed twice with PBST, single infected cells were visualized with DAB substrate (Immunologic # BS04-110) and counted automatically by an ImmunoSpot series 5 UV analyzer (CTL Europe GmbH), as previously described for the forming-focus unit assays developed to study HCV in culture^61,62^. Example of the immunostaining experiments output (images) can be seen in Figures 1d and 2c.

### Treatment assays

Antivirals were obtained from Acme Bioscience and reconstituted in DMSO. For each drug tested, cytotoxicity assays were performed in the different cell types studied, using the Cell Titer AQueous One Solution Cell Proliferation Assay (MTS assay, Promega). The 50% cytotoxic concentration (CC_50_, μM) was calculated by regression analysis. In brief, cells were seeded in 96-well plates and drug dilutions were added the next day, in triplicates. At 72-hr post-infection, the MTS reagent was added, and absorbance was measured after 1 hour incubation at 37°C. Results of treated well were normalized to non-treated controls and the percentage of cell viability was plotted against the Log_10_ of drug concentration, followed by non-linear regression (Y=Bottom + (Top-Bottom)/(1+10^((LogCC_50_−X)*HillSlope)) using GraphPad Prism 8. All treatments shown were conducted at non-cytotoxic concentrations (defined as cell viability ≥80%).

To explore the overall antiviral activity of drug candidates (drug potency), cells were seeded in 96-well plates and the next day, 50 μL of virus (at a specific MOI that led to a robust infection of all non-treated control wells, depending on the cell type) and 50 μL of drug (different concentrations as indicated) were added and the cells were incubated for 72 hr. After the 72-hr incubation, plates were processed for SARS-CoV-2 S-protein immunostaining as described above. The analysis was performed by counting the number of antigen-positive cells which were then normalized to non-treated controls (after subtraction of the background levels obtained from non-infected wells), and EC_50_ (effective concentration 50%) values were obtained after non-linear regression, as stated above.

## Acknowledgements

We thank Bjarne Ørskov Lindhardt (Hvidovre Hospital) and Carsten Geisler (University of Copenhagen) for their support of these studies, and Anna-Louise Sørensen, Susanne Ruszczycka and Louise Barny Christensen, Hvidovre Hospital, for technical support.

